# Quantifying the advantage of multimodal data fusion for survival prediction in cancer patients

**DOI:** 10.1101/2024.01.08.574756

**Authors:** Nikolaos Nikolaou, Domingo Salazar, Harish RaviPrakash, Miguel Gonçalves, Rob Mulla, Nikolay Burlutskiy, Natasha Markuzon, Etai Jacob

**Affiliations:** Oncology Data Science, Oncology R&D, AstraZeneca, Cambridge, UK; Department of Physics & Astronomy, University College London, London, UK; Oncology Data Science, Oncology R&D, AstraZeneca Waltham, MA, USA

**Keywords:** bioinformatics, machine learning, multi-omics integration, multimodal data fusion, late fusion, cancer, survival modeling, TCGA, algorithm evaluation

## Abstract

The last decade has seen an unprecedented advance in technologies at the level of high-throughput molecular assays and image capturing and analysis, as well as clinical phenotyping and digitization of patient data. For decades, genotyping (identification of genomic alterations), the casual anchor in biological processes, has been an essential component in interrogating disease progression and a guiding step in clinical decision making. Indeed, survival rates in patients tested with next-generation sequencing have been found to be significantly higher in those who received a genome-guided therapy than in those who did not. Nevertheless, DNA is only a small part of the complex pathophysiology of cancer development and progression. To assess a more complete picture, researchers have been using data taken from multiple modalities, such as transcripts, proteins, metabolites, and epigenetic factors, that are routinely captured for many patients. Multimodal machine learning offers the potential to leverage information across different bioinformatics modalities to improve predictions of patient outcome. Identifying a multiomics data fusion strategy that clearly demonstrates an improved performance over unimodal approaches is challenging, primarily due to increased dimensionality and other factors, such as small sample sizes and the sparsity and heterogeneity of data. Here we present a flexible pipeline for systematically exploring and comparing multiple multimodal fusion strategies. Using multiple independent data sets from The Cancer Genome Atlas, we developed a late fusion strategy that consistently outperformed unimodal models, clearly demonstrating the advantage of a multimodal fusion model.

## 1. Background

Improving survival predictions (such as overall survival [OS] and progression-free survival [PFS]) in cancer patients is a crucial step in the effort to achieve biological insights and assist clinicians in making more informed clinical decisions. Recent advances in high-throughput sequencing technologies and other molecular assays (such as genomic, transcriptomic, epigenomic, and proteomic methods) have provided a breadth of independent measurements from patients.

Comprehensive integrated analysis of multiomics data can be used to discover the complex mechanisms underlying cancer development and progression. Training predictive models using information from multiple sources can lead to improved model predictions, and many examples, including biological [1–3] and nonbiological [4–6] applications, are reported in the literature. Different data modalities can provide complementary information about patient outcome, including OS. When modalities are correlated, they can help to reduce the variance in these predictions by producing more robust models, which is especially useful when working with data with a low signal-to-noise ratio or a high degree of missingness [5, 7, 8].

Combining data from multiple modalities is challenging, however, and the optimal way to achieve it is largely problem specific [9–13]. Beyond the additional computational and memory burden incurred by increasing the dimensions of the feature space (i.e., biomarkers that constitute model inputs, or independent variables), introducing additional data modalities for a given patient sample also has statistical implications. Because the sample size (number of patients) is fixed, increasing the size of the feature space leads to an increased risk of overfitting and thus the need for regularization. Yet another complication is data heterogeneity. Different modalities might consist of different data types, such as imaging, time-series, text, and tabular data, and often require specific preprocessing and expertise in analysis, modeling, and interpretation. The need for extensive and coherent comparisons across the various multimodal methods is raised repeatedly in several review studies [1, 3, 14–16].

Multimodal data fusion in “omics” data sets usually suffers from low ratios of sample size to feature space. In addition, most individual features are irrelevant or only weakly relevant to the outcome (low signal-to-noise ratio). Some modalities suffer from sparsity of the signal (e.g., mutations) or high degrees of missingness (clinical data), whereas others require batch normalization (gene expression). In addition, the presence of intermodality and intramodality correlations is high [17–20]. These challenges downgrade the potential of multimodal data fusion to add value for biological applications by increasing the likelihood of multimodal-based model overfitting.

The existing literature on data fusion methods using multiomics data for the prediction of OS in cancer patients, although quite extensive [1, 3, 14–16], is characterized by several shortcomings. These include greater emphasis on unsupervised feature reduction methods, linear predictive modeling for survival prediction, modality fusion approaches that ignore the aforementioned properties commonly characterizing omics data, and lack of standardized evaluation approaches that would allow comparison across different methods. The need for extensive and coherent comparisons across the various multimodal methods is a common theme of several reviews [1, 3, 14–16]. Here we summarize the four points we identified as key components in the construction and evaluation of data fusion strategies: dimensionality reduction, survival models, fusion strategies, and evaluation.

### 1.1 Dimensionality reduction

The broad term “dimensionality reduction” encompasses any technique that reduces the size of the feature space. It can refer to “feature selection” techniques that return a subset of the original feature dimensions or to “feature extraction” techniques that return a new, smaller feature set consisting of dimensions that are functions of subsets of the original features. Bioinformatics data sets typically have a very low ratio of samples to number of dimensions, making dimensionality reduction critical for protecting survival models against overfitting. Although typically a relatively small subset of genomic features contributes to survival prediction [21, 22], only a few dimensionality methods have been explored in the context of survival analysis in the literature, which concentrates mostly on unsupervised methods such as Principal component analysis (PCA) or autoencoders [23]. Supervised feature selection methods are limited to the use of univariate Cox proportional hazards (PH) models or training multivariate Cox PH models with Lasso regression (L1-regularization) to impose sparsity [21, 24, 25]. These methods have the advantage of accounting for right-censoring of the data but are quite slow and scale poorly as the feature space increases. They are incapable of accounting for nonlinear correlations with the target (OS time) or for interactions among variables. As such, they limit opportunities for downstream analysis. A range of feature selection methods like Spearman correlation [26] and various information-theoretic approaches [27–31] can address some of these issues.

### 1.2 Survival models

We expected the outcome of a survival model (OS time) to be nonlinearly related to its inputs (biomarkers from the various modalities). Yet the literature largely relies on linear Cox PH models [22, 24, 32]. More recently, deep learning models have also been examined in a few relevant works [33–36], but several nonlinear alternatives, like gradient boosting [37] or random forests [38], are absent from most comparisons. These methods have demonstrated success in survival modeling [39–42] and, although more flexible than linear models, they are equipped with inductive biases that enforce regularization with much less hyperparameter tuning than deep neural networks. Consequently, gradient boosting, random forest, and heterogeneous ensembles [43–45] typically outperform deep neural networks on tabular data [46–48] like multiomics features. Interestingly, although some of these methods have been widely adopted in other biomedical [49, 50] and non-biomedical [51–53] settings, we were not able to find much information on their use in survival analysis.

### 1.3 Fusion strategies

The success of early fusion in other multimodal settings [6, 54–56] has led to recent approaches for modeling predictions of OS in cancer patients with the use of early and intermediate fusion strategies (i.e., data-level fusion). These settings differ from the typical bioinformatics setting, the most important differentiating factor being the number of available data points relative to the input dimensions. For instance, the training set used by Jain et al. [56] contains 1.8–6 billion data points (for the different modalities), whereas The Cancer Genome Atlas (TCGA) data set involves sample sizes on the order of 10–10^3^, depending on cancer type. In the former multimodal application, the feature space is on the order of 10^3^, whereas for TCGA the feature space is on the order of 10^5^. Thus, in the case of multimodal models trained on TCGA [57], late fusion methods (i.e., prediction-level fusion) present an opportunity to outperform early fusion approaches, due to increased resistance to overfitting [16, 58], ease of addressing data heterogeneity, and the ability to more naturally weigh each modality based on its informativeness of OS without being affected by the highly imbalanced dimensionalities across modalities [3, 16].

### 1.4 Evaluation

Many multimodal models for OS prediction that have been reported in the literature suffer from compromised evaluation practices. Some works report validation or even training set results without making use of held-out test data reserved for evaluation. Many fail to account for the considerable uncertainty arising from different training-test set splits of the data. Most lack comparisons spanning different modality combinations or different data fusion approaches. Many even lack direct comparisons against unimodal approaches. Finally, some works propose multimodal approaches in name only, reporting the learning algorithm data from multiple modalities (early fusion) that produced a unimodal model (which used features from only one modality). As a result, most works do not give a straight answer to the questions of whether modalities should be combined and, if so, which ones should be used and how they should be combined.

## 2. Methods

In this work we address these gaps by introducing the AZ-AI multimodal pipeline for the extensive exploration of multimodal fusion strategies that includes several under-explored approaches. We have developed a framework for the extensive exploration of multimodal fusion strategies that allows for rigorous evaluation of multimodal fusion approaches against one another and against unimodal models. We demonstrate the advantage of multimodal data fusion for the cancer patient OS endpoint and outline individual contributions of different modalities. Applying our framework to the TCGA data set, we identify a flexible, ensemble-based, late fusion strategy that consistently takes advantage of additional modalities across independent patient subpopulations for improving OS prediction.

### 2.1 OS prediction

We trained models on the task of predicting OS in patient data from TCGA. In this context, OS measures the length of time (in days) from the date of diagnosis to a patient’s death by any cause. Monitoring OS requires longer follow-up times than other survival endpoints and is affected by deaths due to noncancer causes. Nevertheless, OS is one of the most common endpoints used in practice because there is minimal ambiguity in defining an OS event [59, 60].

As with any survival modeling task, the influence of censoring [61, 62] can complicate the analysis and introduce bias. Although the majority of survival modeling methods listed in Table 1 account for right-censoring, most dimensionality reduction methods presented in Table 2 do not. We used concordance index (C-index) [63] as the most frequently used evaluation metric for the goodness-of-fit of survival models. The C-index is a measure of rank correlation between predicted risk scores and observed survival times. A C-index of 0.5 corresponds to a model whose ranking of predicted patient survival times is no better than random, whereas a C-index of 1 corresponds to a model whose ranking of predicted survival times perfectly matches that of true survival times.

**Table 1.** Cancer types included in TCGA.

| Abbreviation | Cancer type | No. of samples |
| --- | --- | --- |
| BRCA | Breast invasive carcinoma | 1,017 |
| UCEC | Uterine corpus endometrial carcinoma | 513 |
| LGG | Brain lower grade glioma | 509 |
| HNSC | Head and neck squamous-cell carcinoma | 499 |
| PRAD | Prostate adenocarcinoma | 493 |
| LUAD | Lung adenocarcinoma | 491 |
| THCA | Thyroid carcinoma | 489 |
| LUSC | Squamous-cell lung cancer | 470 |
| SKCM | Skin cutaneous melanoma | 450 |
| BLCA | Bladder urothelial carcinoma | 406 |
| STAD | Stomach adenocarcinoma | 402 |
| COAD | Colon adenocarcinoma | 398 |
| KIRC | Kidney renal clear-cell carcinoma | 367 |
| LIHC | Liver hepatocellular carcinoma | 357 |
| CESC | Cervical squamous-cell carcinoma and endocervical adenocarcinoma | 286 |
| KIRP | Kidney renal papillary cell carcinoma | 277 |
| SARC | Sarcoma | 234 |
| OV | Ovarian serous cystadenocarcinoma | 207 |
| ESCA | Esophageal carcinoma | 183 |
| PCPG | Pheochromocytoma and paraganglioma | 178 |
| PAAD | Pancreatic adenocarcinoma | 168 |
| GBM | Glioblastoma multiforme | 155 |
| READ | Rectal adenocarcinoma | 141 |
| TGCT | Testicular germ cell tumors | 128 |
| THYM | Thymoma | 118 |
| LAML | Acute myeloid leukemia | 110 |
| MESO | Mesothelioma | 81 |
| UVM | Uveal melanoma | 80 |
| ACC | Adrenocortical carcinoma | 79 |
| KICH | Kidney chromophobe | 65 |
| UCS | Uterine carcinosarcoma | 57 |
| DLBC | Lymphoid neoplasm diffuse large B-cell lymphoma | 37 |
| CHOL | Cholangiocarcinoma | 36 |
| PANCANCER | All of the above cancer types | 9,481 |

**Table 2.** Modalities included in this work and their preprocessing.

| Modality abbreviation | General description & preprocessing notes | No. of features |
| --- | --- | --- |
| CLIN | Clinical and demographic features<br>Included here: age, gender, race, cancer stage<br>Cancer stage is omitted in early- and late-stage models. | 4 |
| EXP | RNA expression of protein-coding genes (gene expression)<br>Differential expression in normal tissues<br>Selected 1,500 most variable across all indications<br>Selected 1,000 most variable per indication | 1,000 (per individual indication)<br>1,130 (NSCLC)<br>1,500 (PANCANCER) |
| MUT | DNA mutations (binarized; 0: wild type, 1: mutation)<br>Included only those with \$\$VEP_[impact] = 'high'\$\$<br>Selected 1,500 most variable across all indications<br>Selected 1,000 most variable per indication | 1,000 (per individual indication)<br>1,292 (NSCLC) |
| METH | DNA methylation probes (beta values)<br>Retained only those features with <80% missing values<br>Retained only those probes found in CpG islands, within 1,500 base pairs upstream of the transcriptional start site (Chaudhury et al. 2018)<br>Included only differentially methylated for selected genes in EXP<br>Selected 25,000 most variable across all indications<br>Selected 1,000 most variable per indication | 1,000 (per individual indication)<br>1,660 (NSCLC) |
| MIRNA | microRNA expression<br>Retained only those features with <80% missing values | 867 |
| RPPA | Reverse-phase protein array (protein expression) | 198 |
| LNCRNA | Long non-coding RNA expression<br>Extracted from mRNA expression, according to the list in Li et al. [77]<br>Retained only those features with <80% missing values | 1,952 |

mRNA, messenger RNA.

### 2.2 Data

The TCGA project (2006–2018) [57] molecularly characterized more than 20,000 primary cancer and matched normal samples spanning 33 cancer types. From these samples the project produced multiple types of omics data (genomic, epigenomic, transcriptomic, and proteomic), which are listed along with any corresponding demographic, clinical, and imaging data for each sample. Table 1 lists the 33 cancer types in TCGA, along with the number of data points for each, after excluding patients with follow-up times of <1 day. Table 2 lists the modalities included in the experiments along with any preprocessing the samples underwent.

### 2.3 Pipeline overview

Figure 1 provides a summary of the AZ-AI multimodal pipeline. After specifying the subset of interest (cancer type[s] and stage[s], modalities to be included), the data are split into a training set and a test set, the former to be used for feature selection and training models, and the latter used exclusively for the final evaluation. The user specifies the desired train-test split ratio and (optionally) any variable(s) by which the split is to be stratified. The data then undergo optional preprocessing, such as normalizing continuous features or imputing missing data. The transformations applied are learned on the training data and then applied to both training and test data. For example, we performed median imputation by using the training set’s median on both the training and the test sets. Next, the (optional) dimensionality reduction step is employed (see 2.3.1, Dimensionality Reduction), again based on the training set. The chosen survival models are then trained on the training set. A user-specified fraction of the training data is (optionally) used as a validation set to determine the weights of the ensemble (see 2.3.2, Survival Modeling) and of each modality in late fusion multimodal models (see Late Fusion Strategy in figure 2). During model training, fivefold cross-validation is applied to identify the optimal hyperparameter setup under a grid search. The final model is then trained on the full training set and evaluated on the held-out test set. We repeated the entire process multiple times (“runs”) on different training-validation-test splits and computed the average C-index for the test set and the 95% confidence intervals (CIs) for each modality combination (see 2.3.5, Evaluation). The different steps of the pipeline can be executed jointly or per individual modality, giving rise to different multimodal data fusion strategies (see 2.3.3, Fusion Strategies). The user can select multiple cancer types and train a combined model or separate ones for each. Finally, there is the added option to report aggregate permutation feature importances [64, 65] across all runs, affording some degree of model interpretability.

**Fig. 1.**
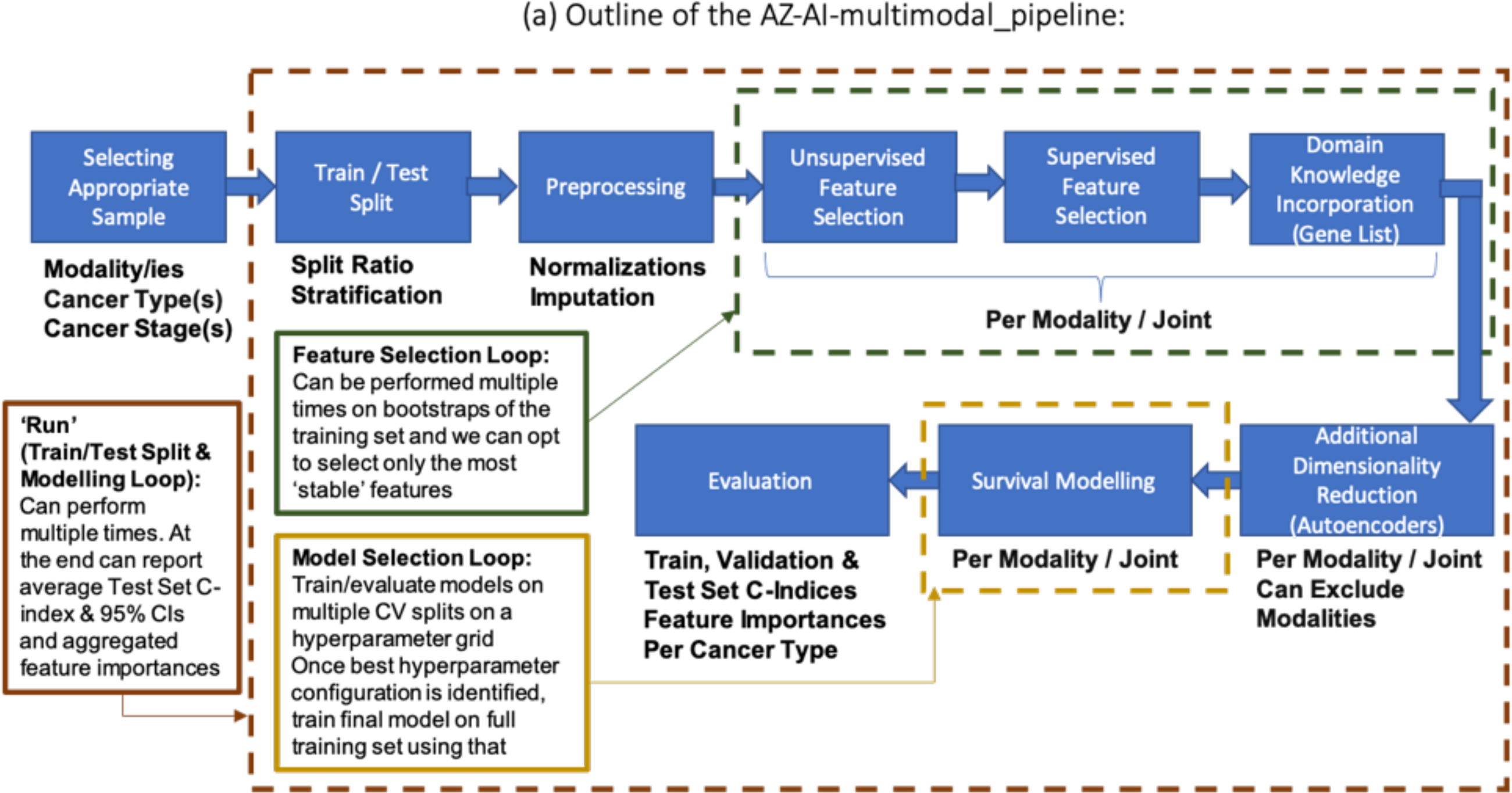
Overview of the AZ-AI multimodal pipeline. (A) Brief outline of the pipeline’s main steps and functionalities. Figure 2 shows a classification of multimodal fusion strategies. The pipeline allows for any of these, depending on which of its execution steps are run “per modality” or “jointly.”

**Fig. 2.**
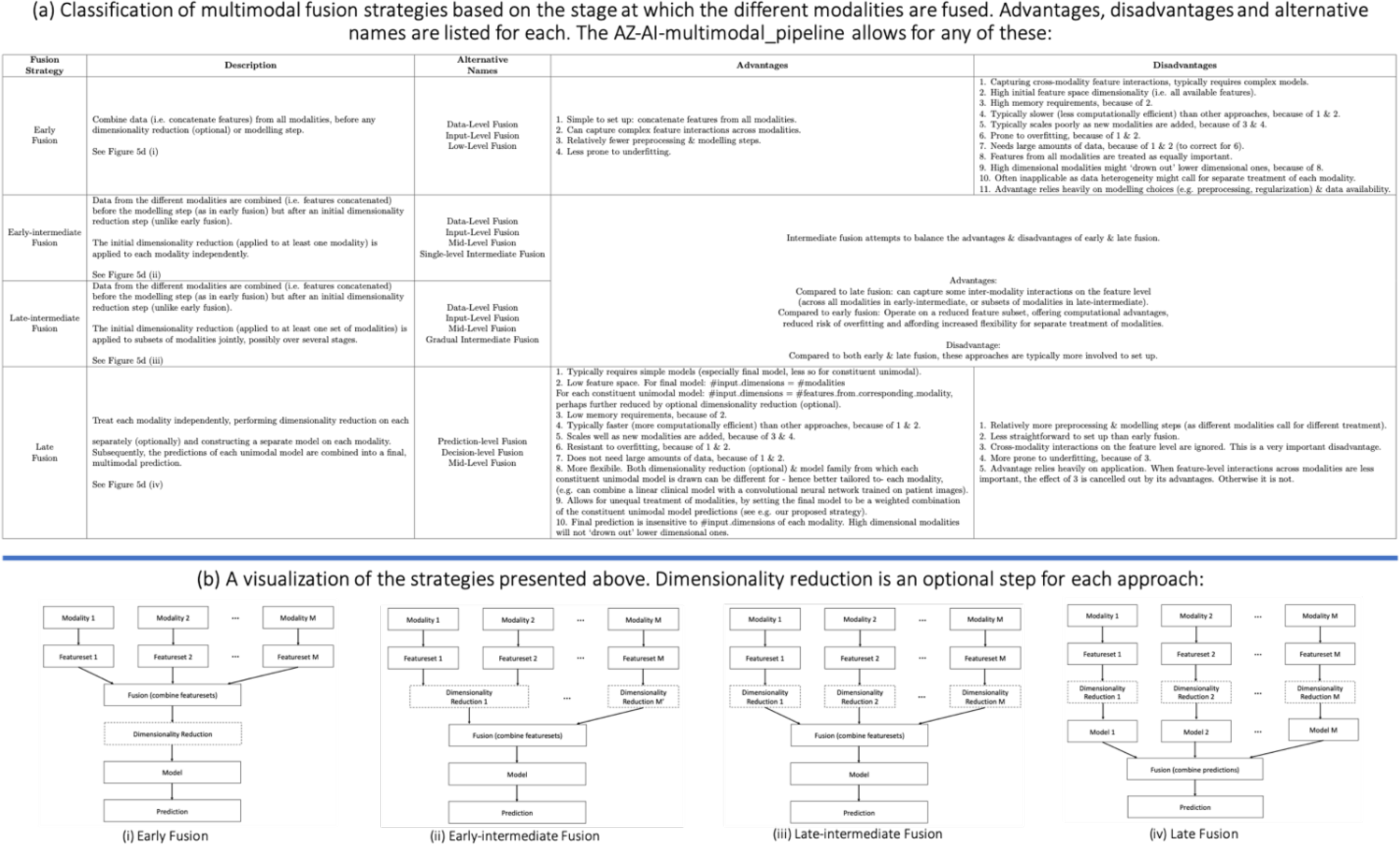
Summary of late, early, and intermediate multimodal data fusion strategies. (A) Description of strategies and their advantages, disadvantages, and alternative names used in the literature. (B) Visual explanation.

#### 2.3.1 Dimensionality reduction

Table 3 list the various dimensionality reduction methods explored. These include both feature selection methods (which return a subset of the original features) and feature extraction methods (which transform the original features), as well as both linear and nonlinear methods, univariate and multivariate methods, and others. Combinations are also possible. For instance, we can first select a subset of the original features by filtering out the least variant ones and then discard features that are highly correlated with other selected ones (both steps unsupervised). We can then train autoencoders to obtain a latent representation of the data (also unsupervised) and, finally, select features that are informative of OS by training univariate Cox PH models.

**Table 3.** Dimensionality reduction methods included in the pipeline.

| Dimensionality reduction method | Subset of original dimensions | Nonlinear (target) | Supervised | Accounts for feature interactions | Accounts for censoring | Incorporates domain knowledge |
| --- | --- | --- | --- | --- | --- | --- |
| Filter correlated features (Pearson) | Y | N | N | Y<sup>a</sup> | N | N |
| Filter low-value variance | Y | N | N | N | N | N |
| Linear correlation (Pearson) | Y | N | Y | N | N | N |
| Monotonic correlation (Spearman) | Y | Y<sup>b</sup> | Y | N | N | N |
| Univariate linear Cox PH models | Y | N | Y | N | Y | N |
| MIM | Y | Y | Y | N | N | N |
| JMI | Y | Y | Y | Y<sup>c</sup> | N | N |
| CMIM | Y | Y | Y | Y | N | N |
| Pathways | N<sup>d</sup> | Y | N | N | N | Y |
| Relevant gene lists | Y | Y | N<sup>e</sup> | N | N | Y |
| Autoencoders | N | Y | N | Y<sup>f</sup> | N | N |

<sup>a</sup>Pairwise interactions, linear, not with regard to target.
<sup>b</sup>Monotonic.
<sup>c</sup>Pairwise interactions, nonlinear, with regard to target.
<sup>d</sup>Biologically meaningful.
<sup>e</sup>Presumed relevant for overall survival.
<sup>f</sup>Dependent on architecture, not with regard to target.
CMIM, conditional mutual information; JMI, joint mutual information; MIM, mutual information maximization; PH, proportional hazards.

There is also the option to specify lists of genes of interest to include for certain modalities. An exhaustive exploration of these approaches and their combinations is beyond the scope of this work, but the provided code can be used to expand on what is presented here. We included the option to limit the risk of overfitting for feature selection approaches by selecting only the most stable features. This can be achieved by applying the feature selection method on *k* bootstraps of the training set and retaining only those features that are selected in at least *k_min_* of the bootstraps by the feature selection method. A bootstrap is generated by uniformly sampling with replacement *N_train_* data points from the original training set of size *N_train_*.

#### 2.3.2 Survival modeling

Table 4 shows the various survival model families included in the pipeline. “ENS” denotes a heterogeneous ensemble of models from all other families listed.

**Table 4.** Survival modeling methods included in the pipeline.

| Survival model method | Nonlinear | Description | Implementation |
| --- | --- | --- | --- |
| CPH-L2 | N | Cox PH model with Ridge (L2-norm) regularization | Scikit-survival [54] |
| CPH-EN | N | Cox PH model with Elastic Net (combined L1-norm and L2-norm) regularization | Scikit-survival [54] |
| RSF | Y | Random survival forest model [25, 26] | Scikit-survival [54] |
| CPH-GB | Y | Gradient boosted [1, 15] Cox PH loss with regression trees as base learner. The partial likelihood of the PH model is optimized as described in Raschka [71]. | Scikit-survival [54] |
| CLS-GB | Y | Gradient boosting [1, 15] with component-wise least squares as base learner [21] | Scikit-survival [54] |
| DS | Y | Deep neural network trained under a Cox PH loss (DeepSurv) [31] | PyCox [34] |
| ENS | Y<sup>a</sup> | Heterogeneous ensemble of any combination of the above models weighted by their respective validation set performances, per Equation 1 | Custom |

<sup>a</sup>Nonlinear, provided at least ONE base model with non-zero weight is nonlinear.
PH, proportional hazards.

Let us denote with *f_i_* the *i*th constituent survival model in the ensemble (trained on the training set). We denote with *f_i_*(***x***) the prediction of model *f_i_* on a data point (patient) with feature vector ***x***. To obtain the ensemble’s prediction *f_ENS_*(***x***) on the same data point, the predictions of each base model (survival risk scores) are first normalized to lie within [0, 1], and then the normalized predictions of all models on the same datapoint ***x*** are linearly combined via weights *w_i_* by:

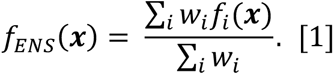

The weight *w*_i_ of each model is determined based on its performance (C-index, *C*_*i*_) on either the training set or on a separate validation set by:

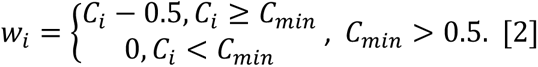

To obtain the results shown, we followed the latter approach. Once the weights *w_i_* were determined on the validation set, retraining the base learners *f_i_* on the combined training and validation set tended to further improve performance, as measured on the held-out test set. We suggest following this approach unless the sample size is too low, in which case we advise using the training set to determine the weights.

Equation 2 weighs the prediction of each model proportionally to its edge over the random survival model performance (*C* = 0.5). Weighted ensembles of this form are common in the literature [66, 67]. Moreover, we added a minimal performance *C_min_* > 0.5 required to include the model in the ensemble [43]. This confers some robustness to the final prediction; poorly performing models are excluded because in small data sets a perceived better-than-random performance can easily be due to chance. The optimal value of *C_min_* is problem specific, but we found that any *C_min_* > 0.5 can improve the ensemble’s performance.

#### 2.3.3 Fusion strategies

A common categorization of multimodal data integration methods is based on when the various modalities are combined and gives rise to three broad classes of fusion strategies: early, late, and intermediate [3, 10, 11, 16, 68]. The relative effectiveness of each depends heavily on the prediction task and the amount of data available [3, 16]. Figure 2 describes each approach and provides a visual explanation along with their respective advantages and disadvantages, as well as alternative names used in the literature.

#### 2.3.4 Proposed late-fusion method

Our proposed late fusion strategy is summarized in Figure 3. It mimics the ensemble construction technique presented in 2.3.2, Survival Modeling. Each modality contributes a model *f_j_*, which itself can be an ensemble of the form of Equation 1. The individual modality predictions are then aggregated using Equation 1. As before, the models *f_j_* are trained on the training set and the weights *w_j_* are determined based on the validation set performance by Equation 2. The predictions *fj*(***x***) can be interpreted as the unimodal risk scores for a patient with feature vector ***x*** and the final ensemble prediction *f_FUSED_*(***x***) as the multimodal risk score. The normalized weight *w_j_*/*∑_j_ w_j_* of the *j*th modality can be interpreted as its relative importance in determining OS in the given setting.

**Fig. 3.**
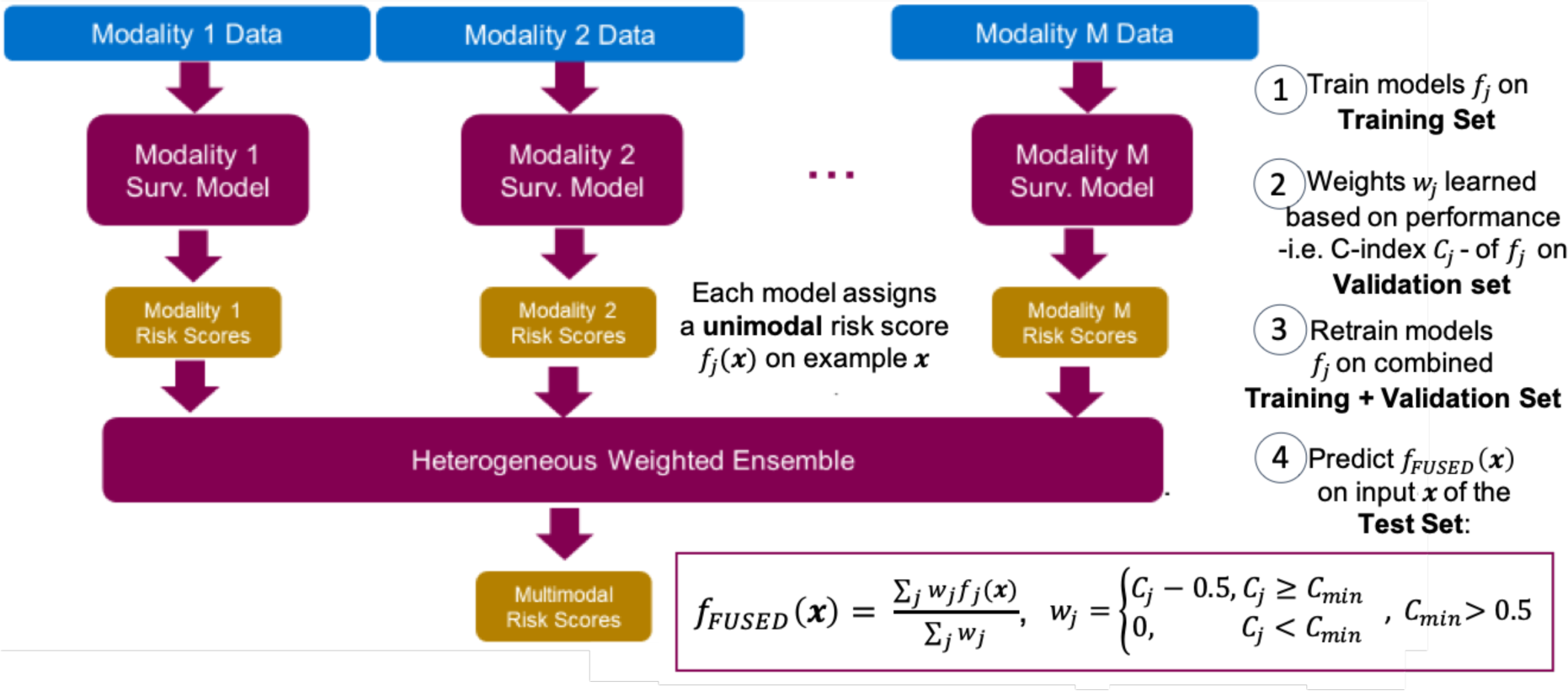
Proposed heterogeneous weighted ensemble-based late fusion strategy.

#### 2.3.5 Evaluation

With very few exceptions [32, 33, 69], studies in the existing literature use a simple evaluation scheme consisting of performing a single train-test split of the data. Assuming best practices are followed and no data leakage is taking place, this is a sensible practice for evaluating a single model on the given train-test split. Some studies [32, 33] go a step further and repeat the model generation process multiple times, reporting average performance and, in the case of Chai et al. [33], a measure of variance of the performance of final models yielded on the same train-test split under their proposed methodologies. This is a sensible way of evaluating the modeling process on the given train-test split.

These evaluation practices, however, ignore the variability resulting from the data split itself, which can be very large, especially in small data sets. One implication is that any results reported (the produced models and their respective performance) are tied to a specific train-test split. This limits reproducibility and renders comparisons across studies meaningless. In the multimodal setting, this practice also hinders comparisons across modality combinations. Train-test split stratification is typically performed on one or more clinical features, but the resulting split can vary greatly in the distribution of nonclinical features. Different splits can therefore favor different modalities. This variability gives rise to the often contradictory results reported in the literature.

Our goal was to answer the question, “Should we combine modalities, and if so, which ones and how?” To claim that one combination of modalities has an advantage over another, or even that there is indeed benefit in combining modalities, it is necessary to evaluate the modeling process across train-test splits under different modality combinations. We thus propose what is known in the machine learning literature as “repeated holdout validation” [70–73] as a more appropriate choice of validating the results in this setting. For each modality combination, we trained multiple models on different training sets and evaluated each on the corresponding held-out test set, reporting average performance and 95% CIs.

#### 2.3.6 Experimental setup

For all results presented here, unless stated otherwise, we constructed multimodal models using the strategy described in 2.3.4, Proposed Late Fusion Method. We first split the data set into 80% training and 20% test data. For numerical features, we performed median imputation and then selected the top 25 features on each modality by Spearman correlation with OS time. For mutations in particular, we provided a prespecified list of the top 25 genes associated with each indication as identified by AstraZeneca’s Biological Insights Knowledge Graph (BIKG) [78]. BIKG combines relevant data for drug development from public and internal data sources. On the selected feature subset, 80% of the available training data are then used to train survival models and perform hyperparameter optimization by using fivefold cross-validation. The remaining 20% of the training set was used as a validation set. We then evaluated the models by predicting the OS of the test data (20% of total data). We repeated the process 10 times on different train-validation test splits and computed the average test set C-index and 95% CIs for each modality combination. The variance across runs can be very high, especially when modeling smaller subsets of TCGA (see Table 2 for sample sizes of each subpopulation). Therefore, we repeated the train-test split multiple times when comparing the relative importance of each individual modality or modality combination. We rounded results to the third decimal digit. For both unimodal and multimodal models, unless stated otherwise, we report the results of ensembles of survival models of all families shown in Table 4 except DS (excluded due to its relatively slow hyperparameter optimization), combined under Equation 1. We set *C_min_* = 0.53 for Equation 2.

## 3. Results

### 3.1. Multimodal data integration pipeline

The AZ-AI multimodal pipeline, developed in the context of this work by the AstraZeneca Oncology Data Science Team, is a Python library for multimodal feature integration and survival prediction. It can be used to preprocess and reduce the dimensionality of tabular data sets (unimodal or multimodal) and to train and evaluate survival models. Its functionalities include several preprocessing and imputation options, flexibility regarding when to integrate modalities (Figure 2), a range of feature reduction approaches (Table 3) and survival modeling methods (Table 4), and rigorous evaluation (as described in 2.3, Pipeline Overview), including the option to report the models’ feature importance. An outline of the pipeline is provided in Figure 1.

The library can be used to replicate and extend the results presented in this paper. It has already been successfully used for constructing state-of-the-art early integration multimodal models combining clinical and radiological features for OS prediction in lung cancer patients in a forthcoming paper by Patwardhan et al. (manuscript in preparation). The setting examined in this paper is quite different from that of those investigators. As a result, different methods proved more successful in each application. In the work by Patwardhan et al., the data consist of only two modalities, the number of total features was on the order of 10^2^–10^3^, and the number of data points was on the order of 10^3^, all being complete cases (no need for data imputation). This paper examines data sets of four to seven modalities, with missing entries and the total number of features on the order of 10^3^–10^5^ and the number of data points being 10–10^3^, depending on cancer type (approaching 10^4^ only for pancancer models). As a result, the risk of overfitting here is considerably higher. In the setting of Padwardhan et al., early and intermediate fusion strategies and nonlinear and nonmonotonic feature selection methods (mutual information) worked best. In our setting, late fusion strategies and linear or monotonic feature selection methods (Pearson and Spearman correlation) outperformed the other approaches. In both cases, ensemble survival models outperformed single models. This comparison demonstrates that different approaches to multimodal fusion are better suited to different settings, simpler feature selection methods and late fusion are more suitable for situations in which the risk of overfitting is high, forming ensembles of multiple survival models is always beneficial, and our AZ-AI multimodal pipeline is flexible enough to provide a good multimodal fusion solution in different settings.

### 3.2 Results on using late fusion in TCGA cancer patients

Throughout this paper, we report the average test set C-index and 95% CI across 10 runs. The experimental setup, proposed late fusion strategy, and details on the evaluation and the data set are all described in 2.2, Data. Unless stated otherwise, the results correspond to the feature selection method being AAA and the survival model being a heterogeneous ensemble of all models listed in Table 4.

#### 3.2.1 Performance of multimodal versus unimodal models across different cancer types

For each cancer type in TCGA, we calculated the advantage (henceforth referred to as the “delta”) of the multimodal approach over unimodal models as the difference in average C-index between the multimodal model and the best unimodal model:

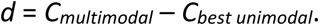

Figure 4 shows the *d* across all 33 TCGA cancer types. In 25 of 33 cancer types (76% of the independent data sets), the deltas were positive, supporting the hypothesis that late fusion multimodal models generally outperform unimodal models. This result is statistically significant (*P* = 0.001) under a Wilcoxon signed-rank test against the null hypothesis of no multimodal advantage (*d* = 0). Aligned with our expectations of the impact of the sample size, the multimodal advantage *d* positively correlated with the training set size. The associated Pearson correlation *r*(*d*, sample size) = 0.361 is statistically significant (*P* = 0.039). Finally, *d* also positively correlates with the performance of the best unimodal model. In other words, the better the best unimodal model, the larger the advantage of using it in multimodal models. This might seem counterintuitive, but it is largely due to our late fusion strategy, which, because it combines all unimodal models into a multimodal model, requires at least one strong base model to achieve good performance. That said, this finding might also indicate other mechanisms worth exploring, such as the presence of strong cross-modality correlations; if this is the case, then when one modality is predictive of OS, others would be as well, and combining them into multimodal models would further improve results. The associated Pearson correlation adjusted by weighting data sets by sample size *r_adj_*(*d*, *C_best unimodal_*) = 0.365 is statistically significant (*P* = 0.029).

**Fig. 4.**
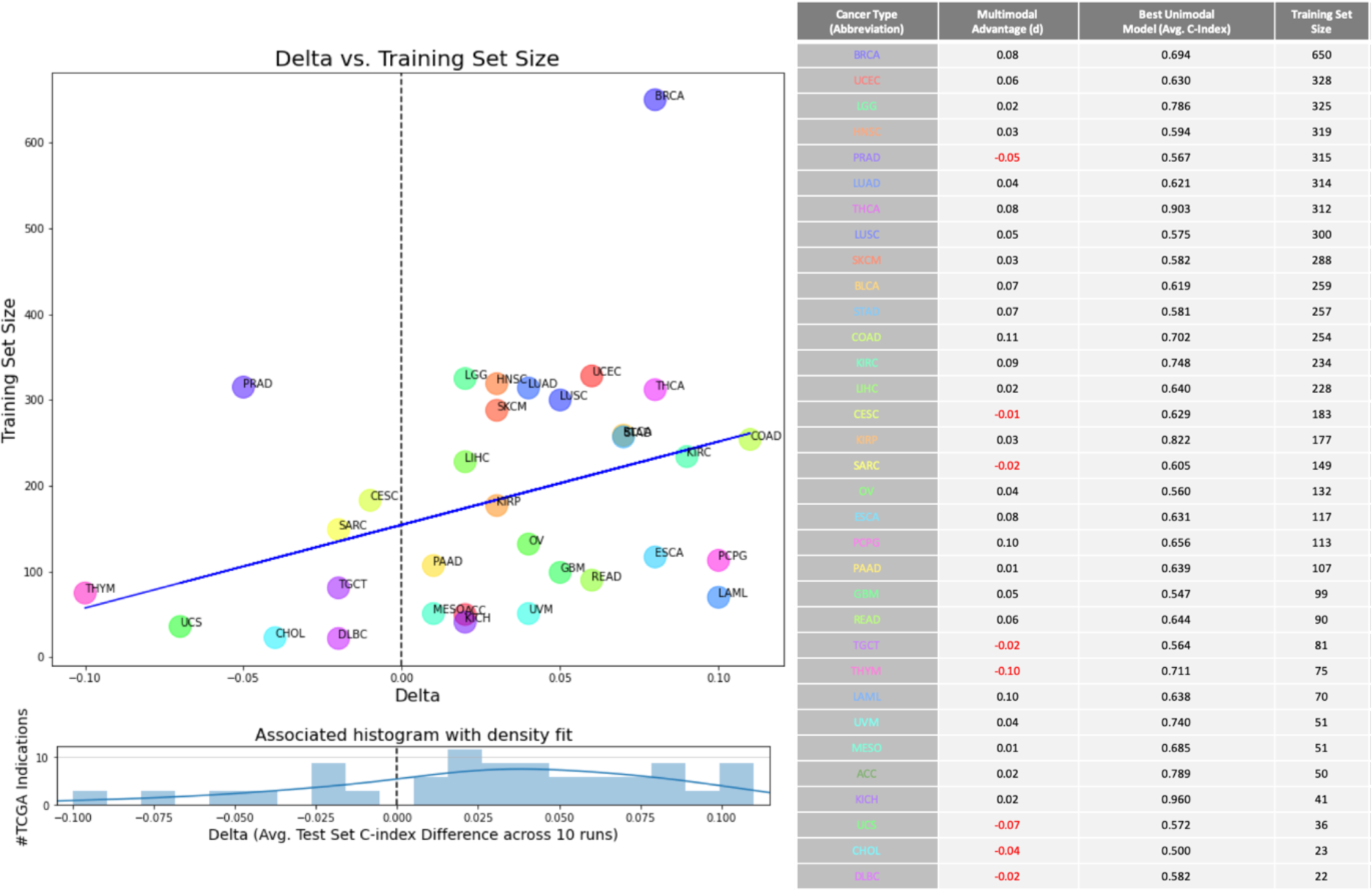
[Right] Table showing the advantage *d* of multimodal versus best unimodal model in terms of average C-index across all 33 TCGA cancer types. The table also lists the average C-index of the best unimodal model and the size of the training set per cancer type (entries listed in decreasing order). [Left, top] Multimodal advantage *d* versus training set size. The two quantities are positively correlated (blue trend line added for emphasis). Advantage *d* and training set size are also positively correlated. [Left, bottom] Associated histogram of advantage *d* with density fit shown. Dashed line denotes no advantage (*d* = 0). Multimodal models dominate best unimodal ones in 25 of the 33 indications (*d* > 0).

Summarizing the above results, we found that multimodal models trained under our proposed late fusion strategy tended to outperform unimodal models across TCGA cancer types. We also found that the advantage of multimodal models increased with sample size and with the performance of unimodal models.

#### 3.2.2 Comparing multimodal with unimodal models, NSCLC, and other cancer types

Within the TCGA data, non–small-cell lung carcinoma (NSCLC) was the most well-represented cancer type, with more than 1,000 patients having a diagnosis of lung adenocarcinoma (LUAD) or lung squamous-cell carcinoma (LUSC). We therefore discuss the NSCLC results in more detail. In addition, we included more modalities than we did for other TCGA cancer types, allowing a more comprehensive analysis.

In Figure 5, we compare the performance of unimodal models from each available modality against multimodal models trained under our proposed late fusion strategy (FUSED). Figure 5A shows results for models trained jointly on NSCLC patients, and Figures 5B and 5C show results on its two subtypes (LUAD and LUSC) separately (A joint treatment of both subtypes of NSCLC is common in the literature [35, 74]). Univariate Cox PH models were used for feature selection. Similar results obtained for all indications included in the TCGA data set are provided in the Additional File section 1.

**Fig. 5.**
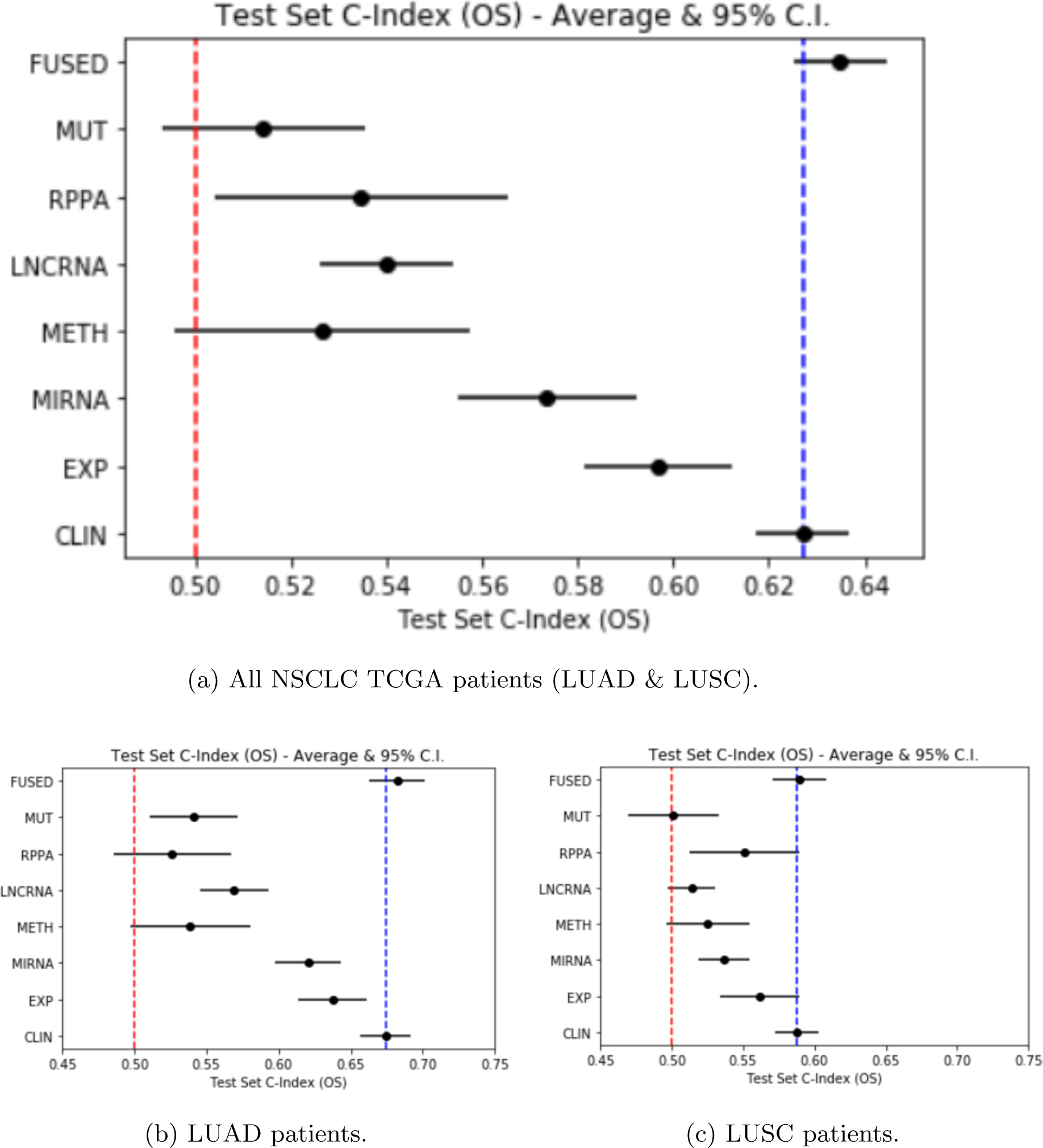
Performance of multimodal models (FUSED) versus unimodal models of each modality for (A) all NSCLC patients, (B) LUAD patients only, and (C) LUSC patients only. See Additional File section 1 for other TCGA cancer types. Average test set C-index and 95% CI across 10 runs are reported. The red dashed line denotes random prediction performance (C-index = 0.5). The blue dashed line denotes the average C-index of the best individual modality (here: clinical features). Multimodal models outperformed all unimodal models on average and had lower variance. * Figs. 4 and 5 are based on different feature selection methods. (Fig. 4 required many more executions and therefore we used a faster feature selection method).

Regarding individual modalities, Figure 5 suggests that each carried some signal for predicting OS, since unimodal models of any modality (with the exception of mutations, in the case of LUSC patients) attained an average C-index of >0.5 on new data. Not all individual modalities were equally predictive of OS. For example, across all NSCLC patients, clinical and demographic features (CLIN) constituted the best individual modality, mainly due to including cancer stage, a strong predictor of OS. They are followed by gene expressions (EXP), with mutations (MUT) the weakest individual modality, possibly because of its binary treatment. The high variance across runs (which justifies our evaluation strategy) permits us to compare only the average performance of each modality.

Comparing unimodal models to the multimodal model, we observed a small advantage in terms of average C-index of the multimodal model over even the best individual modality (here: clinical features). We also found that the variance of the multimodal models was considerably reduced (despite the increased feature space). The results applied to patients of both NSCLC subtypes, although the advantage of multimodal models was reduced, possibly due to the smaller sample size. These observations generally held beyond NSCLC across all TCGA indications (see Additional File section 1).

The relative importance of each modality in the multimodal model (based on the weights calculated on the validation set) correlated well with the performance of each unimodal model (based on average C-index on the test set) (see Additional File). Inspecting the weights also verified that the multimodal model did not reduce to a unimodal model (no weight was equal to 0, meaning all modalities were considered).

#### 3.2.3 Exhaustive comparison of all modality combinations in NSCLC

Figure 6 shows the average test set C-index for all possible modality combinations on NSCLC patients, under all models included in the study. The results suggest that adding more modalities leads to improved survival predictions (in the worst, best, and average cases). The performance improvement appeared to plateau after including the two to three most informative modalities, and adding more beyond this point conferred diminishing benefits. In addition, excluding the best individual modality (here: clinical) from the fused model resulted in a large drop in performance (≤4–7.5%, depending on the subpopulation) between combinations of modalities that did and did not include clinical features. Similar results on all cancer types for modalities EXP, MUT, reverse-phase protein array, and CLIN are included in Additional File section 4.

**Fig 6.**
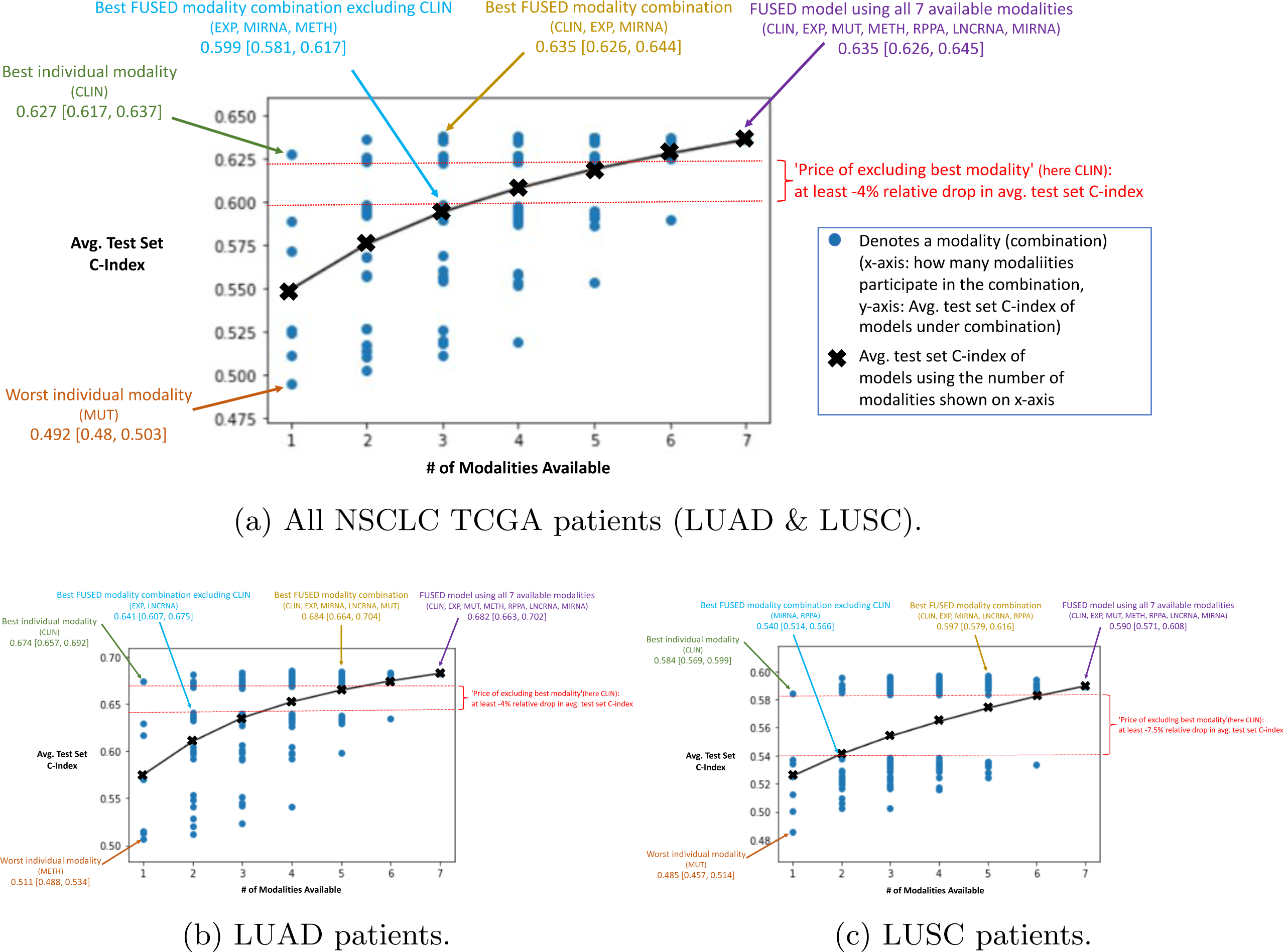
Average test set C-index for each of the 7! possible modality combinations (blue points) on the subset of (A) all NSCLC TCGA patients (LUAD and LUSC), (B) LUAD patients only, and (C) LUSC patients only. The black crosses indicate the average C-index across all multimodal models trained on *k* modalities, where *k* is equal to the number shown in the x-axes (black trend line added for emphasis). On average, the more modalities added, the better the resulting model. The red dashed lines mark the effect of not including the best individual modality in the multimodal fusion. Also shown is the average test set C-index and 95% CI for: (i) the worst individual modality (orange), (ii) the best individual modality (green), (iii) the best modality combination excluding the clinical features (light blue), (iv) the best modality combination (gold), and (v) the multimodal fusion of all seven modalities (purple). We note the diminishing benefit of adding more modalities and the high “price” of excluding the best individual modality (here: clinical features).

Exhaustive modality combination plots such as the one shown in Figure 6 are a useful tool for determining whether adding modalities improves performance and by how much, as well as which modalities should be combined. Such a plot would be computationally intensive to produce for an early fusion strategy, but it can be efficiently obtained under a late fusion strategy, as all underlying unimodal models are already trained.

### 3.3 PANCANCER model results

Figure 7 shows the results of a comparison of the performance of unimodal models and multimodal models trained on the subset of all TCGA patients (PANCANCER). The multimodal model significantly outperformed all unimodal models (*P* < 0.0001), including that of the best-performing individual modality (clinical features), and exhibited decreased variance. For all TCGA stages and cancer types, multimodal (FUSED) models had an average (± standard deviation) test set C-index of 0.785 ± 0.005, whereas the best unimodal (CLIN) had an average test set C-index of 0.763 ± 0.007 across 10 runs. On this larger sample of patients, our proposed late fusion strategy clearly showed benefits over unimodal models. To our knowledge, these are the top performing results attained by TCGA pancancer models [34, 75]. In contrast to individual cancer types (e.g., NSCLC in Fig. 5), aggregating over all 33 indications, we found no noticeable differences on the modality level between early-stage patients (Fig. 5A) and late-stage patients (Fig. 5B).

**Fig. 7.**
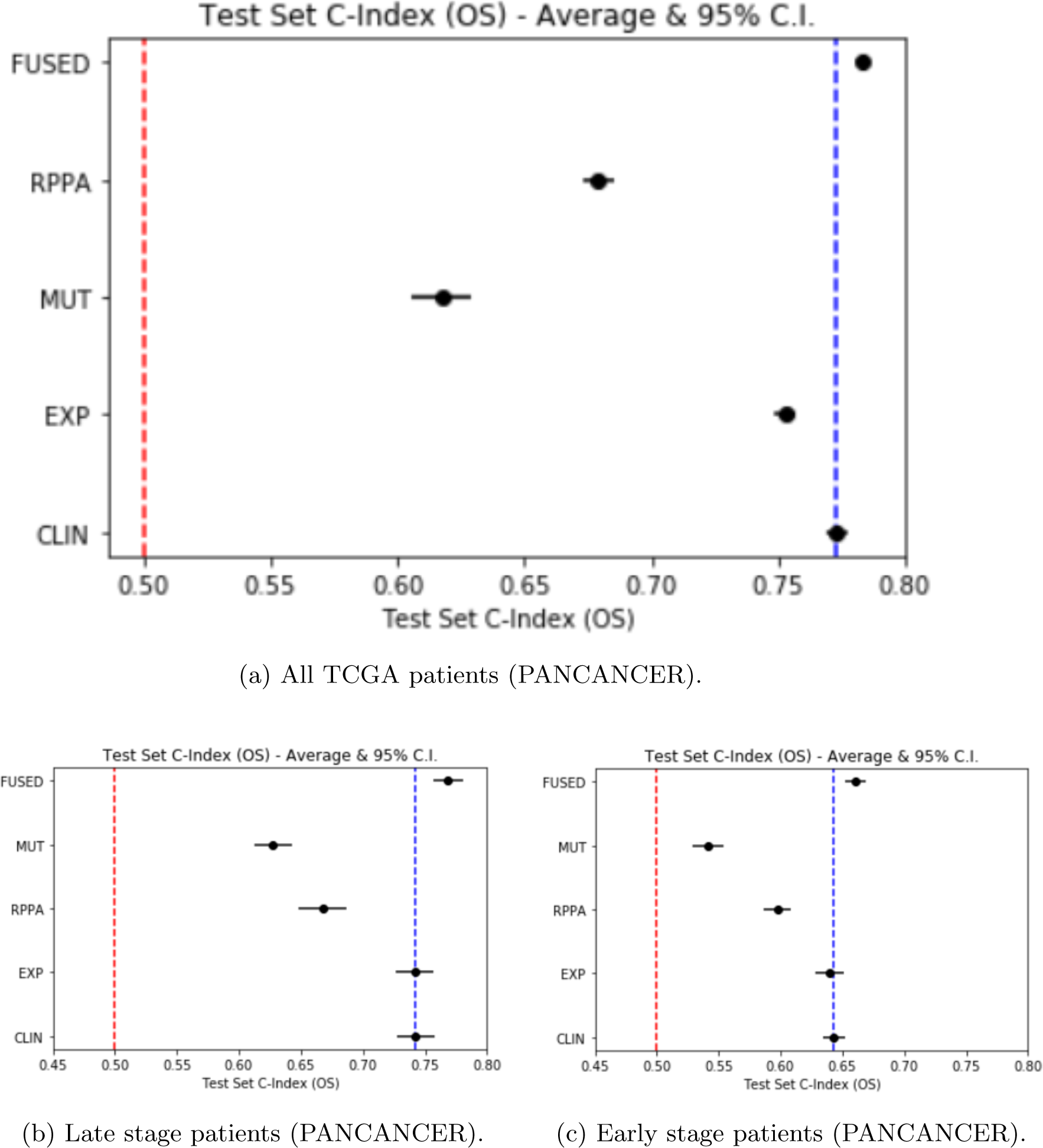
Performance of multimodal models (FUSED) versus unimodal models of each modality trained on TCGA patients of all cancer types (PANCANCER) for patients of (A) all stages, (B) late stages (III and V), and (C) early stages (I, II). Average test set C-index and 95% CIs across 10 runs are reported. The red dashed line denotes random prediction performance (C-index = 0.5). The blue dashed line denotes average C-index of best individual modality (here: clinical features). Multimodal models significantly outperformed all unimodal models and had lower variance.

In these subpopulations, cancer stage (clinical feature) was removed. For both groups, clinical features and gene expressions were tied for best individual modalities for predicting OS. Among the clinical features, cancer type is an important indicator of OS. A likely explanation for gene expression being equally predictive is that, because different types of cancers affect different tissues and different tissues are characterized by distinctive gene expression profiles, gene expression is used by the model as a proxy for cancer type. This is a reminder that information from different modalities can be overlapping. Nevertheless, obtaining the same information from different data sources can decrease the multimodal model’s uncertainty. This is another advantage of multimodal models, evidence for which is shown in these results.

Pancancer models are not uncommon in the multiomics literature [34, 75], as the larger sample size lowers the risk of overfitting (which increases as more modalities are added). The clinical usefulness of pancancer models is limited, however, as they are trained and evaluated on 33 different types of cancers. We included these results primarily to showcase the power of our proposed fusion strategy on a larger data set. However, there might still be benefits in the multimodal setting of training models on a pancancer level to make predictions on individual cancer types. An example is provided in Additional File section 5.

## 4. Discussion

We observed a consistent advantage of multimodal data fusion in these experiments. Multimodal models outperformed the best unimodal models on 24 of all 33 TCGA cancer types (see 3.1, Multimodal Data Integration Pipeline, and Additional File section 1). These models exhibited reduced variance and better average predictive performance, showing clear superiority in larger sample sizes (see 3.1, Multimodal Data Integration Pipeline; 3.2, Multimodal Data Integration Pipeline; and Additional File sections 1 and 2). We found that adding modalities yielded diminishing benefits but did not adversely affect performance (see 3.3, PANCANCER Model Results). Finally, we found that on pancancer models, where the sample size was larger, the multimodal advantage was particularly pronounced (3.3, PANCANCER Model Results). Under our proposed late fusion strategy, we concluded that we should combine any available omics data. We expect larger benefits with larger training sets and more informative individual modalities.

Across the different subpopulations examined, clinical features and gene expressions generally carried a stronger signal for predicting OS than other modalities (e.g., mutations). This could reflect the limitation of our modeling (e.g., binary encoding of mutations), but most likely it is due to the models being inherently more informative. For example, clinical features include variables like “cancer type” and “cancer stage,” which are expected to be strong predictors of OS. We hypothesize that gene expression acts as a proxy for “cancer type” (different tissues have different gene expression profiles) and, possibly to some extent, “cancer stage.” As a result, multimodal models that do not use these modalities tend to be inferior to those that do, and adding more modalities tends to confer only minor advantages (mostly in terms of reduced variance). This might reflect modeling limitations or could be evidence that the various omics modalities are largely redundant given gene expression. More uncorrelated sources of information, like imaging data, could also be beneficial, and early works, including those by Patwardhan et al. (in preparation), Cheerla and Gevaert [34], and Wulczyn et al. [69], suggest that this is the case.

Other interesting findings include the observation that all modalities were used by the resulting multimodal models, and their relative importance roughly agreed with their predictiveness of OS (Additional File section 2). Our initial exploration into differences across cancer stages suggests that the role of protein expressions and mutations in late-stage NSCLC and breast invasive carcinoma (BRCA) patients in determining OS is greater than in early stages (Additional File section 3). Finally, we demonstrated an example of a multimodal pancancer model outperforming models trained on a single cancer type (Additional File section 5).

The pipeline described in 2.3, Pipeline Overview, is in no way exhaustive but is quite extensive and can be easily adapted to include new methods. It would be especially useful for benchmarking and exploring general tendencies (e.g., early vs. late fusion). The proposed fusion strategy, however, does have limitations. As a late fusion method, it cannot capture cross-modality feature interactions. We did not choose it by an exhaustive search over the possibilities offered by the AZ-AI multimodal pipeline but rather for its simplicity, flexibility, and scalability and because it yields consistently good results that demonstrate the added benefit of multimodal data fusion (as suggested by our results). On different data sets (e.g., with larger sample size–to–dimensionality ratio, higher signal-to-noise ratio, or higher degree of cross-modality interactions), we do not necessarily expect it to outperform other approaches, but its qualitative advantages (simplicity, flexibility, and scalability) will still hold.

## 5. Conclusions

Bioinformatics data sets typically consist of various modalities, such as genomic, transcriptomic, epigenomic, proteomic, lifestyle, and phenotypic. Multimodal data fusion offers the potential to improve patient outcomes and gain novel insights by combining information across modalities. Multiple strategies can be used to integrate multimodal data, however, and they are highly problem specific; identifying an appropriate approach for a given setting is challenging. The literature on predicting OS in cancer patients is fragmented and inconsistent, and several promising methods remain underexplored. A consistent and rigorous comparison of multimodal methods against one another and against unimodal models is also missing from the currently available literature. Such comparisons are necessary to determine whether different omics data sets should be combined and if so, which ones should be used and how they should be combined.

To answer these questions, we implemented the AZ-AI multimodal pipeline, which allows us to train, evaluate, and compare a large number of multi-omics methods for survival analysis. We identified a late fusion strategy that consistently demonstrated the advantages of multimodal data fusion of clinical and demographic features, DNA mutations, gene-coding RNA expressions, long non-coding RNA expressions, microRNA expressions, methylations, and protein expressions for the prediction of OS in patients in TCGA. This advantage was consistently demonstrated across individual TCGA cancer types and on pancancer models. Multimodal models exhibited reduced variance and better average predictive performance, showing clear superiority in larger sample sizes. Finally, we found that adding modalities did not harm performance, although the benefits diminished. We therefore conclude that, with our proposed strategy, we should combine different omics data sets.

Clinical and demographic data, along with data on differential gene expression, were found to be the most informative modalities overall. Our initial explorations into differences between early- and late-stage NSCLC and BRCA patients suggest an increased role of protein expressions and mutations in patients with late-stage disease in determining OS. A promising future direction would be to further explore the relative importance of modalities for different patient subpopulations, especially in relation to a given treatment. Beyond the modality level, identifying key biomarkers that drive the prediction would require the use of model interpretability methods such as permutation feature importance (included in the AZ-AI multimodal pipeline) or SHapley Additive exPlanations (SHAP) [76].

Our results, along with those of Patwardhan et al. (in preparation), support the claims of others [3, 16, 58] that late fusion is more successful than early fusion in settings where there is high signal-to-noise ratio and low sample size–to–dimensions ratio, due to a decreased risk of overfitting. A detailed exploration of problem characteristics such as the these and more (e.g., degree of intramodality and intermodality correlations) to optimal fusion strategy, which is possible with the use of the AZ-AI multimodal pipeline, is left for future work.

Finally, it would be interesting to explore more ways of incorporating domain knowledge into the models. Working with pathways and gene sets is a promising direction that could reduce multimodal model dimensionality by leveraging knowledge of the underlying biology and could allow more successful intermediate fusion approaches for more biologically interpretable insights to be drawn from the models.

## Declarations

### Ethics approval and consent to participate

Not applicable.

### Consent for publication

Not applicable.

### Availability of data and materials

The data sets generated during and/or analyzed during the current study are available in the [NAME] repository, [PERSISTENT WEB LINK TO DATASETS].[Reference number]

### Competing Interests

All authors are or were employees of AstraZeneca at the time this work was performed and may have stock ownership, options, or interests in the company.

### Funding

This study was funded by AstraZeneca.

### Authors’ contributions

[TODO: Add]

## Acknowledgments

The authors thank Jacob Gould Ellen, Gustavo Alonso Arango-Argoty, and Paul Metcalfe for their helpful suggestions that improved this manuscript.

