## Supplemental material for "Quantifying the advantage of multimodal data fusion for survival prediction in cancer patients"

**Supplemental Files**

In Section 3.1 of the main paper, we discuss aggregate results comparing the performance of multimodal survival models (FUSED) obtained via our proposed late fusion strategy (Section 2.3 in main paper) against unimodal models for all cancer types included in The Cancer Genome Atlas (TCGA) dataset. The modalities included are clinical and demographics data (CLIN), gene expressions (EXP), mutations (MUT), and protein expressions (RPPA).

Here we present the detailed results for each cancer type. In figures shown here, we present the average test set C-index and 95% confidence interval (CI) across 10 runs for each modality. Spearman correlation with overall survival (OS) time was used for feature selection for all modalities except MUT, where AstraZeneca’s Biological Insights Knowledge Graph (BIKG) [1] was used to identify top 25 mutations associated with each type of cancer. The red dashed line denotes random prediction performance (concordance index [C-index] = 0.5). The blue dashed line denotes the average C-index of the best individual modality for each cancer type. An aggregation of these results is presented in Fig. 1 of the main paper. The size of the training set differs across cancer types (see Fig. 1).

**Fig. S1**

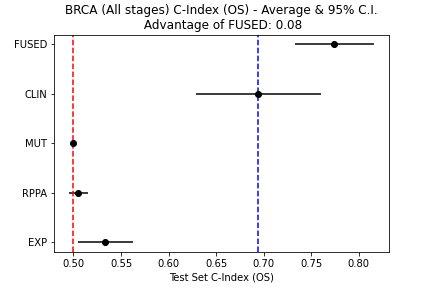

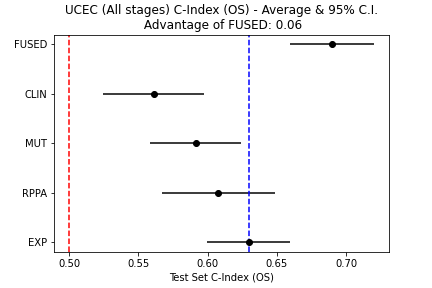

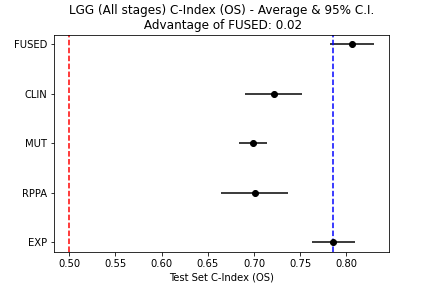

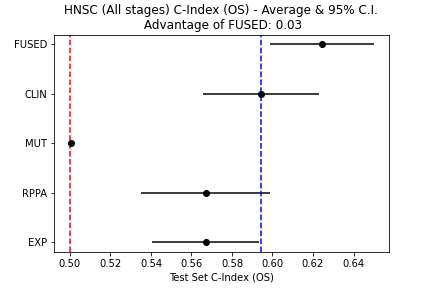

**HSNC**

**LGG**

**UCEC**

**BRCA**

**LUSC**

**THCA**

**LUAD**

**PRAD**

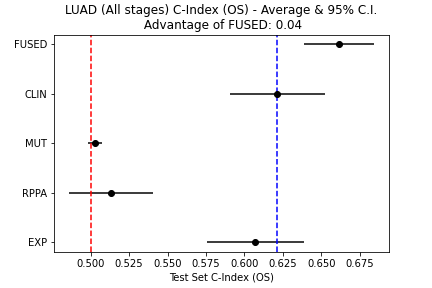

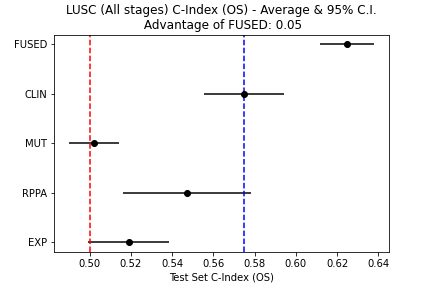

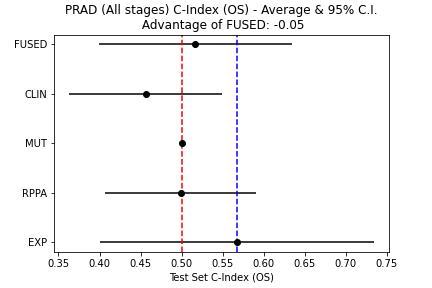

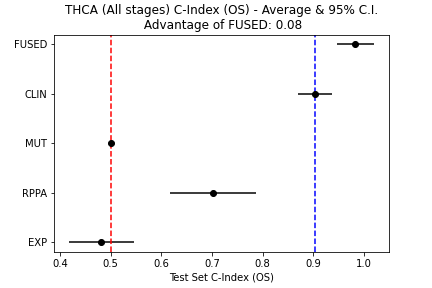

**KIRP**

**CESC**

**LIHC**

**KIRC**

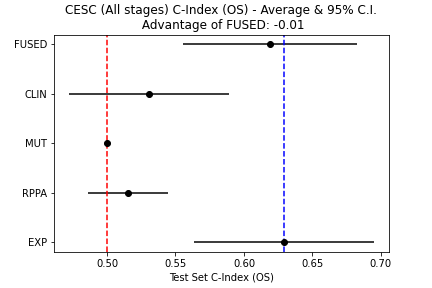

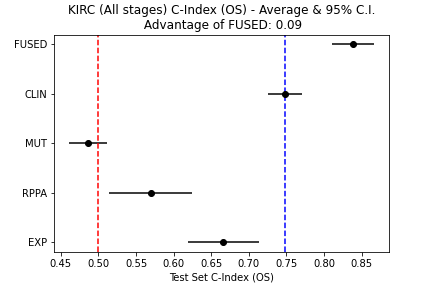

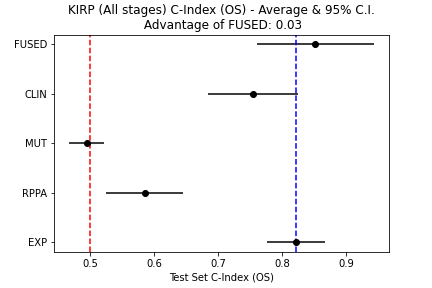

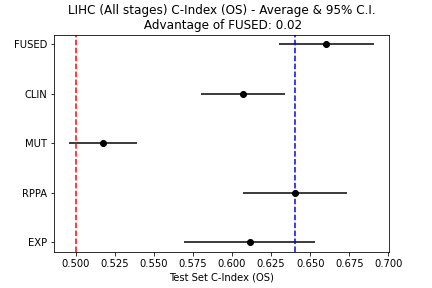

**PCPG**

**ESCA**

**OV**

**SARC**

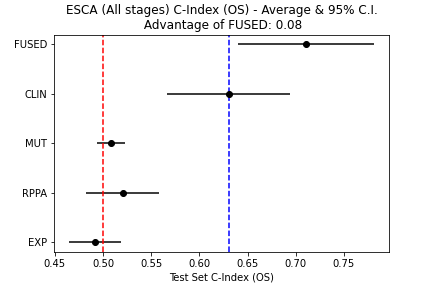

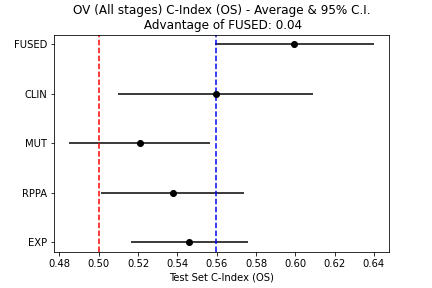

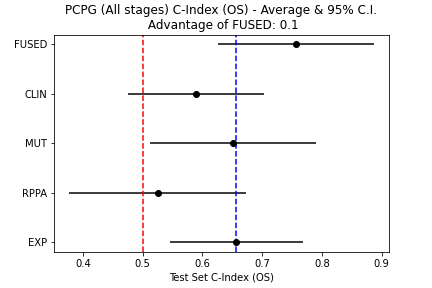

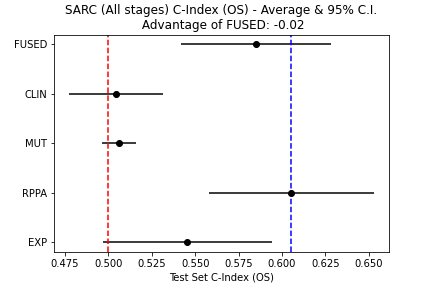

**PCPG**

**ESCA**

**OV**

**SARC**

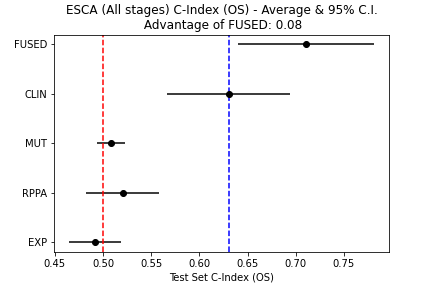

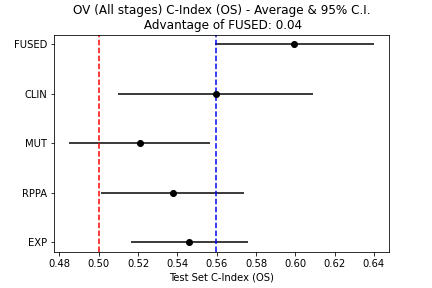

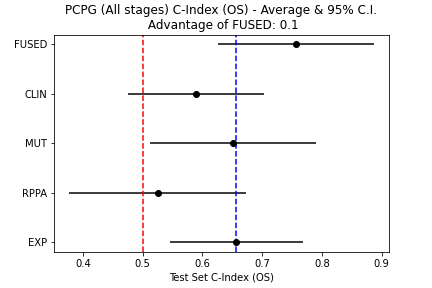

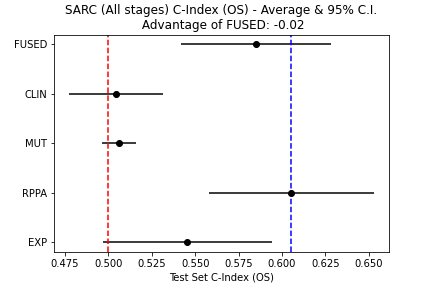

**MESO**

**UVM**

**LAML**

**THYM**

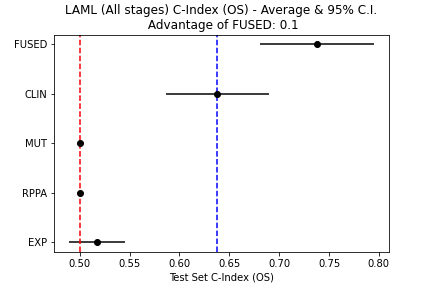

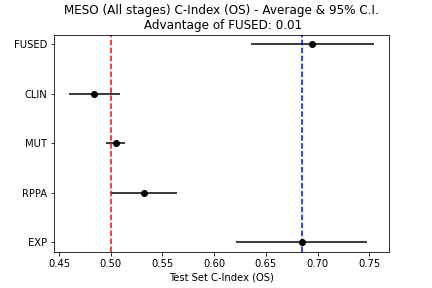

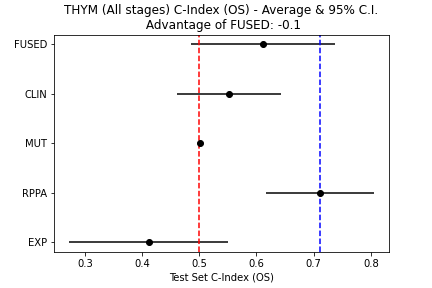

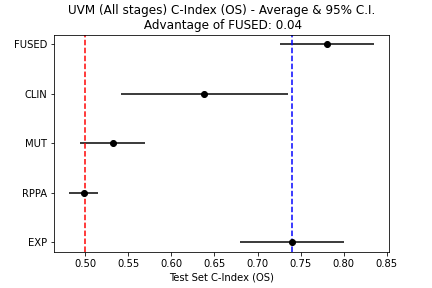

**CHOL**

**UCS**

**KICH**

**ACC**

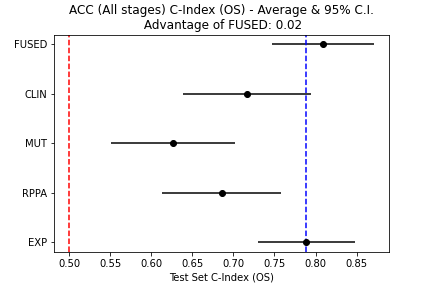

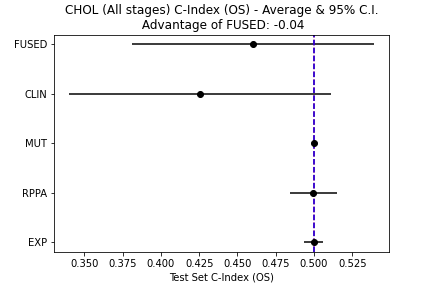

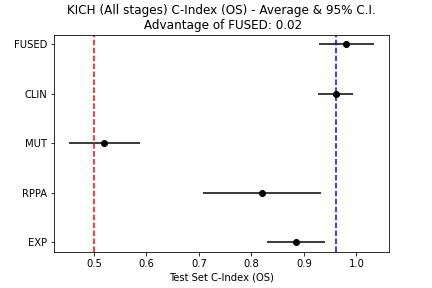

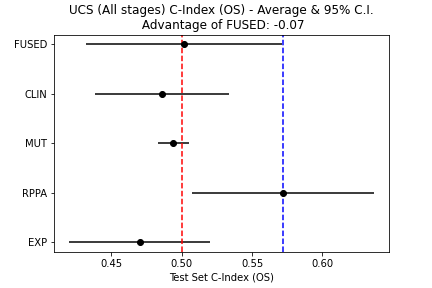

**DLBC**

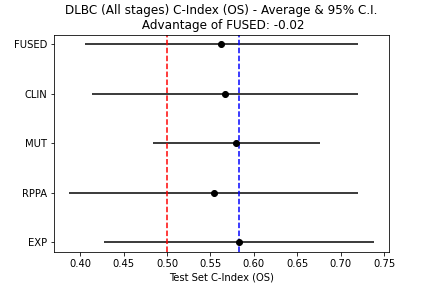

**2	Investigation of relative importance of each modality (NSCLC)**

In Section 3.2 of the main paper, we note that the multimodal (FUSED) model does not reduce to a unimodal one and uses all modalities (no modality has a weight of 0). We also mentioned that the weight assigned by the multimodal model to each modality roughly agrees with that modality’s predictivity of OS.

To demonstrate the above, we show the average (normalized) weight of each modality in the fused model for patients with non–small-cell lung carcinoma (NSCLC) (Table S1). The rank of each modality (1 = most important to 7 = least important) is added for clarity. All modalities are assigned a nonzero weight by the multimodal model. The relative importance of modalities (based on weights calculated on the validation set) generally correlates well with the performance of each unimodal model on the relevant subpopulation’s test set.

**Table S1**

| **Modality** | **Cancer type** | | | | | |
| --- | --- | --- | --- | --- | --- | --- |
|  | **NSCLC (LUAD & LUSC)** | | **LUAD** | | **LUSC** | |
|  | **Average weight** | **Average rank** | **Average weight** | **Average rank** | **Average weight** | **Average rank** |
| CLIN | 0.608666 | 1 | 0.4597 | 1 | 0.511612 | 1 |
| EXP | 0.241698 | 2 | 0.313532 | 2 | 0.079312 | 4 |
| MIRNA | 0.050685 | 5 | 0.095268 | 3 | 0.081792 | 3 |
| METH | 0.005704 | 4 | 0.028218 | 4 | 0.070946 | 5 |
| LNCRNA | 0.018155 | 3 | 0.049727 | 6 | 0.030299 | 7 |
| RPPA | 0.045146 | 6 | 0.009351 | 7 | 0.177833 | 2 |
| MUT | 0.009945 | 7 | 0.044133 | 5 | 0.048205 | 6 |

LUAD, lung adenocarcinoma; LUSC, lung squamous-cell carcinoma.

**3	Differences between early- and late-stage patients (NSCLC)**

Here we explore differences on the modality level between early-stage (I and II) and late-stage (III and IV) NSCLC (lung adenocarcinoma [LUAD] and lung squamous-cell carcinoma [LUSC]) patients. We inspect how the relative performance of unimodal models to predict OS trained on each modality differs across the two subpopulations. Features for each modality were selected using univariate Cox proportional hazards (PH) models.

The results are shown in Fig. S2. Noting that the sample sizes for the two subpopulations differ considerably (there are 756 patients with early-stage and 194 with late-stage NSCLC in TCGA), we observed some interesting differences regarding the potential importance of each modality between early- and late-stage NSCLC patients.

Mutations (MUT) and protein abundances (RPPA) from least important indicators of OS in early-stage patients become more important for late-stage patients. Although clinical and demographic features (CLIN) constitute the most important modality for early-stage patients, they become the least important one for late stages. Gene expression (EXP) becomes the most informative modality in late stages.

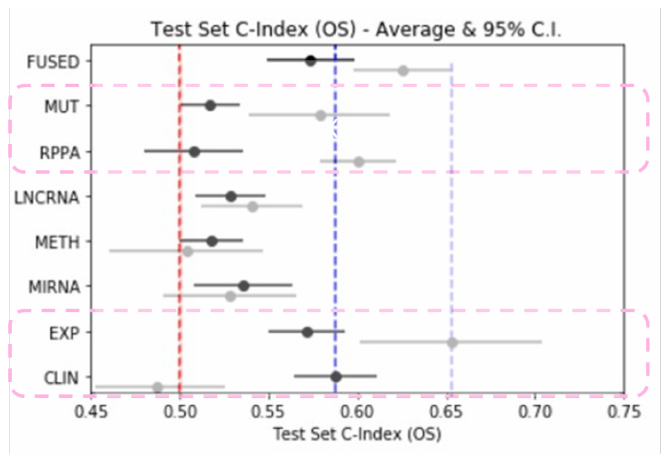

**Fig. S2**	Average test set C-index and 95% CI across 10 runs for each modality for models trained and evaluated on data from patients with early-stage (black) and late-stage (grey) NSCLC. The red dashed line denotes random prediction performance (C-index = 0.5). The blue dashed lines denote the average C-index of the best individual modality for each subpopulation.

**4	Results on each individual TCGA indications (exhaustive comparison)**

In Section 3.3 of the main paper, we present an exhaustive comparison of survival models trained under all possible combinations of seven available modalities for NSCLC patients, summarized in Figure 3 of the main paper. Here we include detailed results for all cancer types included in TCGA for clinical and demographic data (CLIN), gene expression (EXP), mutations (MUT), and protein expression (RPPA).

In Fig. S3, each point corresponds to the average test set C-index for each of the 4! possible modality combinations. The black crosses show the average C-index across all multimodal models trained on *k* modalities, where *k* is equal to the number shown in the x-axes (black trend line added for emphasis). On average, the more modalities added, the better the resulting model.

**Fig. S3**

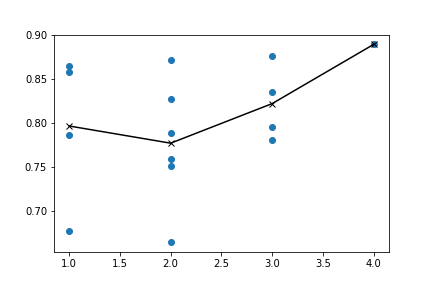

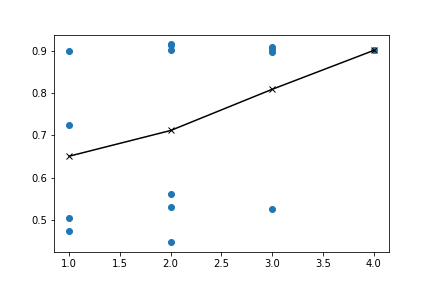

KICH

THCA

LGG

ACC

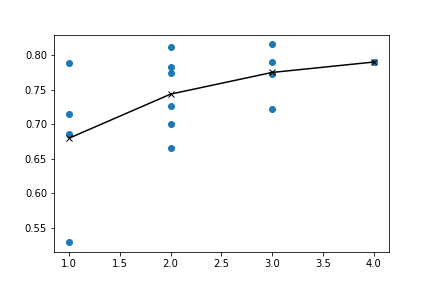

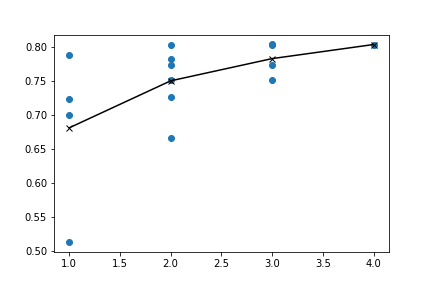

KIRP

UVM

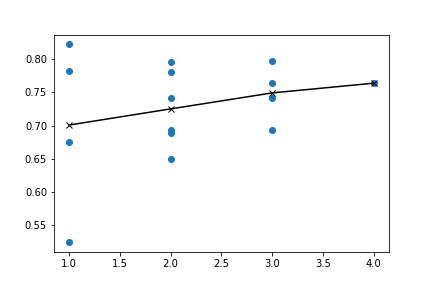

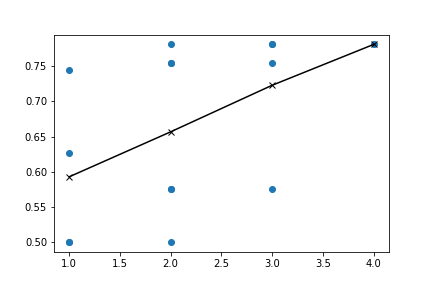

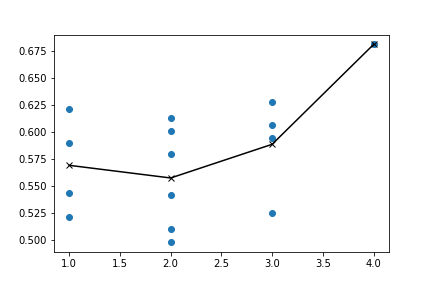

KIRC

COAD

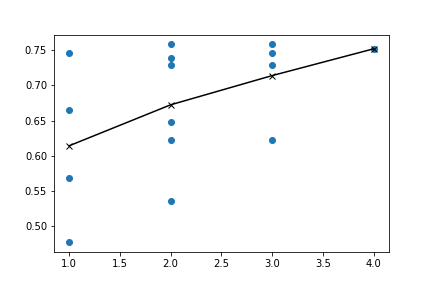

BRCA

PCPG

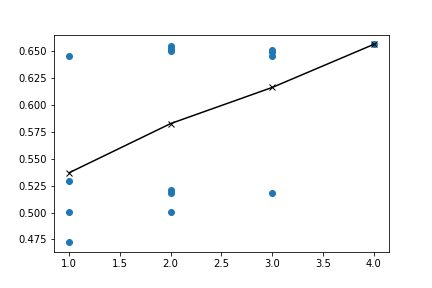

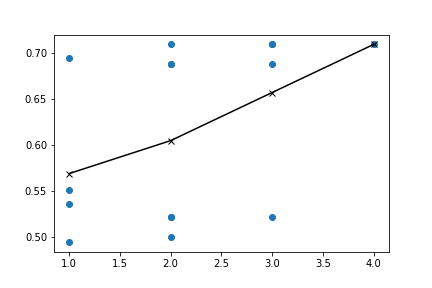

MESO

UCEC

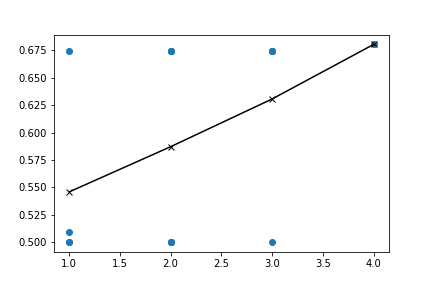

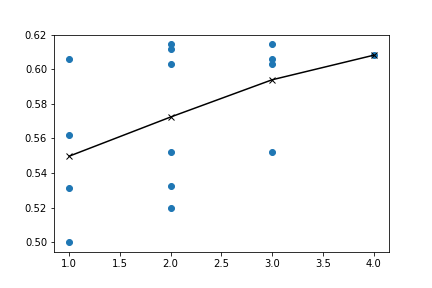

LIHC

LUAD

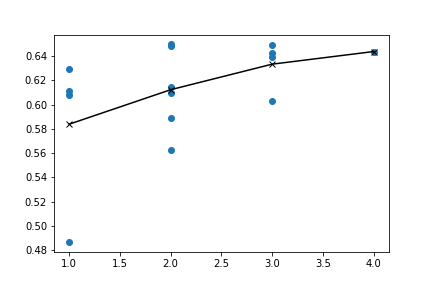

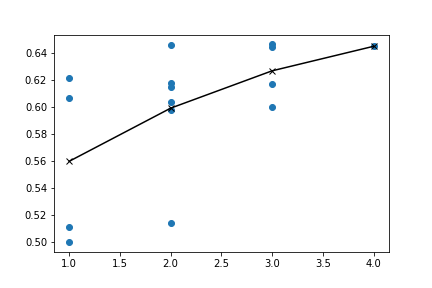

PAAD

LAML

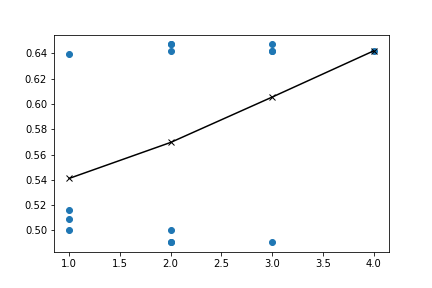

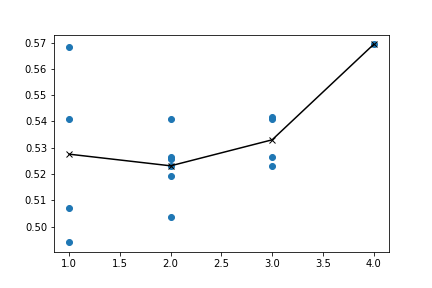

BLCA

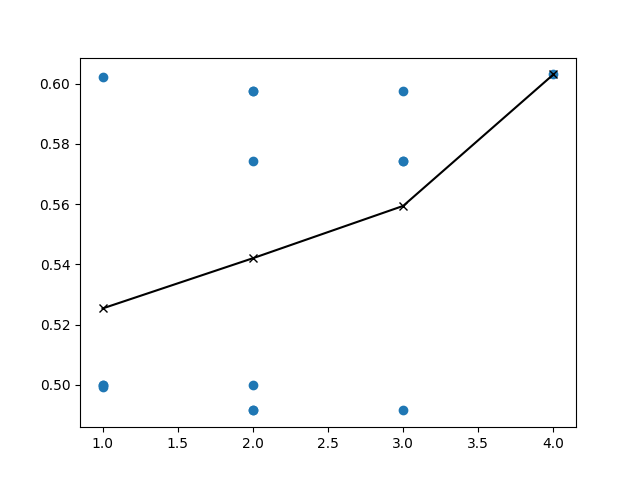

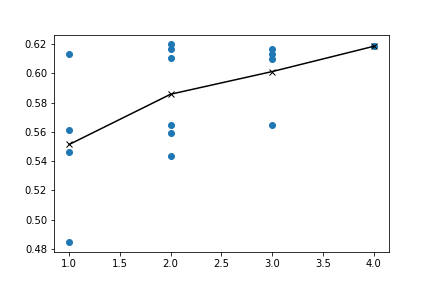

CESC

THYM

SKCM

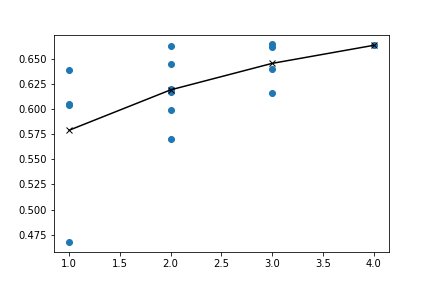

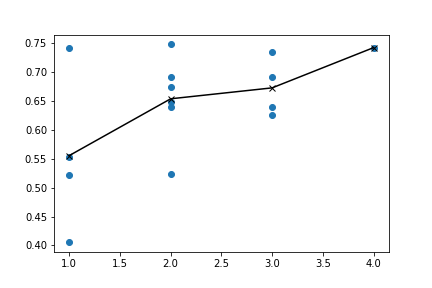

ESCA

HNSC

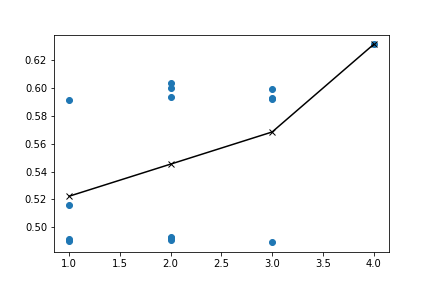

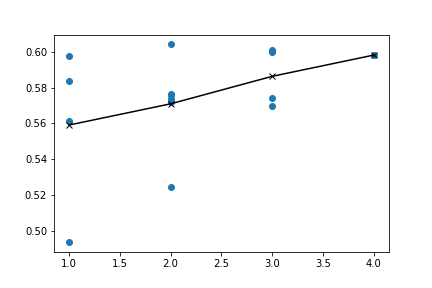

LUSC

READ

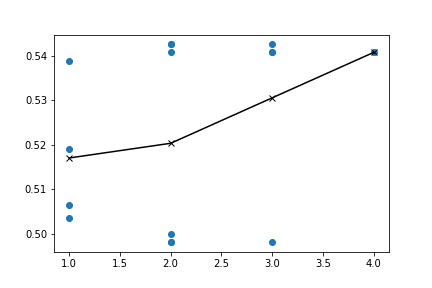

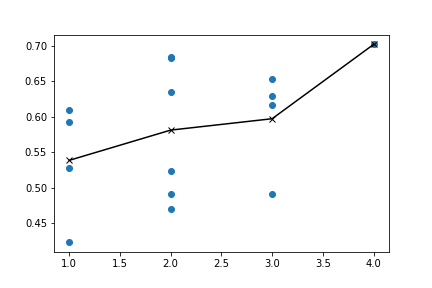

OV

STAD

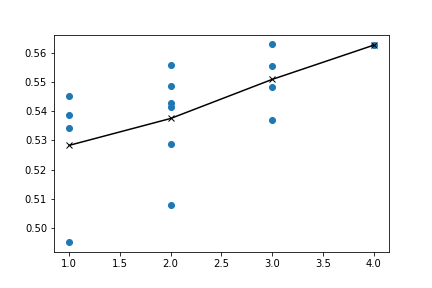

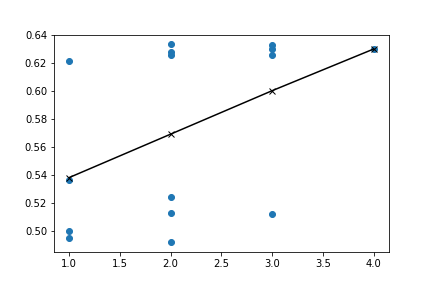

SARC

DLBC

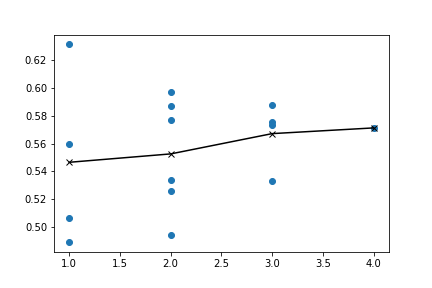

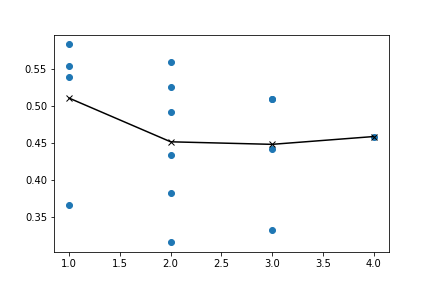

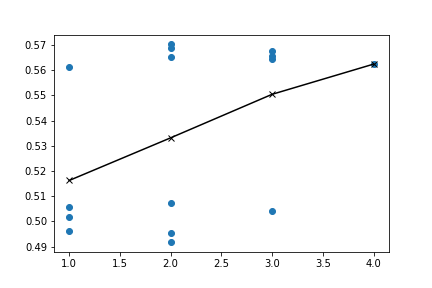

GBM

TGCT

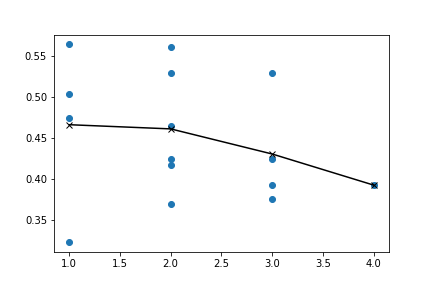

UCS

PRAD

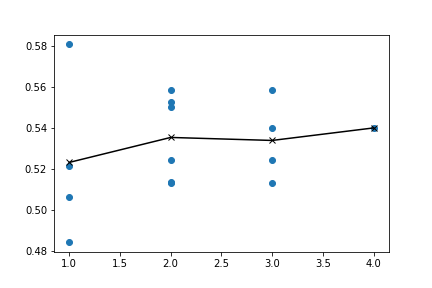

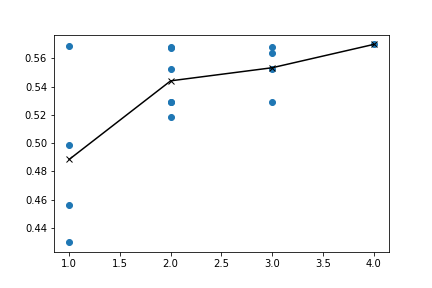

CHOL

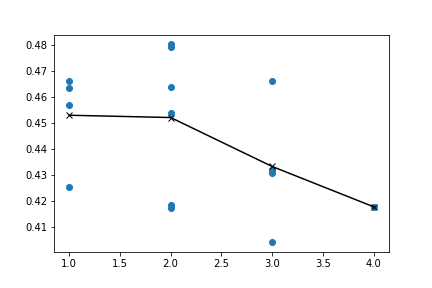

**5	Pancancer models can outperform indication-specific models**

In Section 3.4 of the main paper, we note that there might be benefits in the multimodal setting of training models on a pancancer level to make predictions on individual cancer types. To showcase this, we compared models trained on a pancancer training set of late-stage (III and IV) cancer patients against models trained only on a BRCA late-stage patient training set.

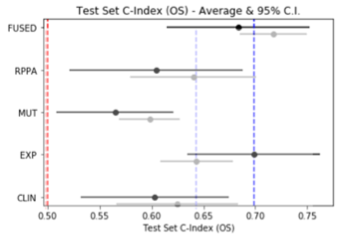
The results are shown in Fig. S4. We see that fused models trained on a pancancer sample of patients outperform the unimodal models as well as the multimodal models trained specifically on BRCA patients. On the level of individual modalities, we find that by training on the pancancer sample, all modalities become better indicators of OS except for gene expression. Protein expressions and mutations showed the biggest improvement. In other words, in this setting (late-stage BRCA patient OS prediction), unimodal models for protein expressions and mutations seem to be particularly benefited by a larger training sample (pancancer), rather than a sample specific to BRCA. Possibly these modalities are the main drivers for the improvement of the respective multimodal model as well. We found
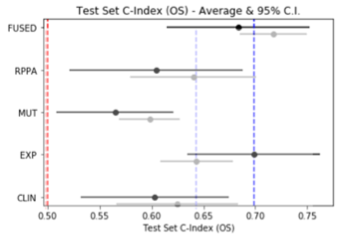
that this was not the case for early-stage BRCA.

**Fig. S4** Average test set C-index and 95% CI across 10 runs for each modality for models trained on late-stage BRCA patients (black) and late-stage TCGA patients (PANCANCER) (gray) and evaluated on late-stage BRCA patients. The red dashed line denotes random prediction performance (C-index = 0.5). The blue dashed lines denote the average C-index of the best individual modality for each subpopulation
